# Nonlethal effects of predation: Presence of insectivorous birds affects the behaviour and level of stress in insects

**DOI:** 10.1101/2022.04.01.486535

**Authors:** Jan Kollross, Jitka Jancuchova-Laskova, Irena Kleckova, Inga Freiberga, Dalibor Kodrik, Katerina Sam

## Abstract

Insect exposure to their predators can affect individuals and community processes, through direct consumption or nonlethal (i.e., nonconsumptive) effects. However, the links between behavioural and physiological responses and stimuli needed for development of the fear are not clear. We therefore subjected the desert locusts (*Schistocerca gregaria*) to three nonlethal treatments, using the great tits (*Parus major*) as a potential predator. The treatments involved: (1) *bird* - presence of a live great tit and its calls, (2) *call* - great tit calls only, (3) *control* - without any treatment. In the first behavioural laboratory experiment, hungry locusts were kept in an experimental cage with a shelter and food on opposite sides of the cage. The duration of hiding and feeding were considered as an indicator of fear responses. In the second laboratory experiment with the same three treatments, levels of the adipokinetic hormone (AKH) were evaluated in the central nervous system (CNS) and haemolymph. In the third experiment in an outdoor aviary, birds were free to fly in larger distances from locusts, before hormone levels were measured as response to bird and control treatments. In the first behavioural experiment, the presence of tits and their call resulted in significantly longer hiding time and significantly shorter feeding time than in the call/control treatments. The proximity of birds and locusts in the laboratory experiment elicited a significant increase in the AKH levels in the CNS and haemolymph as compared to the call/control treatments. In the outdoor experiment, the AKH levels were significantly higher in the CNS of locusts exposed to the bird than to control; no difference was recorded in their haemolymph. We showed that predator exposure quickly affected behavioural responses and physiological processes of locusts. Playback of the avian calls was not an appropriate stimulus to induce stress responses in desert locusts.

---

Insectivorous predators have a tremendous impact on insect communities whose effect cascades further down to plants (e.g., (Hawlena and Schmitz, 2010; Yang and Gratton, 2014). They can affect prey populations by direct consumption (lethal effects or ‘density’ effects) and also through nonlethal effects (trait-mediated effects or interactions). During the nonlethal effects, behavioural compensations for fear of predation occur, leading to alteration of prey properties, their populations and associated communities (Buchanan et al., 2017; Miller et al., 2020). Nonlethal effects are often defined operationally as any predator-caused factor that reduces the population growth via a reduction in survival and/or developmental performance (Schmitz et al., 1997; Nelson et al., 2004; Schmitz, 2007; 2009).

Although most of the existing studies focus on the lethal effects of predators, the nonlethal effects may be even more important to population dynamics (Buchanan et al., 2017; Miller et al., 2020). Indeed, some of the most famous examples of lethal vertebrate predator-prey interactions, are now believed to operate not solely through consumption but also through intimidation which leads to changes in prey’s traits and behaviour (Lind and Cresswell, 2005; Peckarsky et al., 2008; Dröge et al., 2017). Efforts to avoid death make animals to forgo foraging, reduce activity, seek refuges, and exhibit other costly predator-induced defences.

These behavioural responses seem to be uniform both for vertebrates and invertebrates, despite they are more often studied in vertebrates. The presence of invertebrate predators can further lead to earlier maturation at the smaller size when growth rate does not change, or slower growth rate leading to later maturation at the same or smaller size in invertebrates (Abrams, 1995; Abrams and Rowe, 1996).

While each predation event only influences a single prey per unit of time, the risk introduced by the mere presence of a predator could have more widespread effects by causing many prey individuals to significantly alter their foraging behaviour in that same period of time (Cinel et al., 2020). Moreover, trade-offs are probably included in insect prey responding to various threats. For example, presence of invertebrate predators can reduce the resilience of insects to the pesticides (Op de Beeck et al., 2016) and nonlethal effects of predators can thus further affect prey mortality and impact ecosystem functioning. Understanding of such nonlethal effects of insectivorous predators is especially important as insects are the most dominant animal group on Earth (Stork et al., 2015), serve as prey for many different predators and consume up to 70% of total leaf area in some habitats. Although insects have an extraordinary diversity of anti-predator behavioural and physiological responses, predator-induced stress has not been studied extensively in insects (Cinel et al., 2020).

It can be assumed that insects activate their biochemical and physiological defence systems after contact with predators, as it is common for other stressful situations (Cinel and Taylor, 2019) and then change their behaviour accordingly. Such responses are mostly unknown (Kodrík et al., 2015; Cinel et al., 2020) because stress response in the insect body might be difficult to detect and quantify – both behaviourally and physiologically. Stress stimuli represented by the presence of predator induces secretion of the stress hormones from the central nervous system (CNS), specifically from the corpora cardiaca, a small neuroendocrine gland connected with the brain, and hormones further spread by haemolymph to the other body parts (Nässel and Zandawala, 2020). In insects, the anti-stress reactions seem to be regulated predominantly by adipokinetic hormones (AKHs) and octopamine (Cinel et al., 2020). AKHs act as typical anti-stress hormones: they mobilize lipid, carbohydrate, and amino acid proline energy stores (Gade et al., 1997) by stimulating catabolic reactions to gain energy. Simultaneously, AKHs inhibit synthetic reactions; thus, the mobilized energy is used to eliminate imminent stress situations. It has been documented that increase of AKHs in insects depends on stress situations and signals (Kodrik, 2008).

Signals which trigger stress reactions in insects are rarely studied. However, it has been shown that lizard-specific vibrations and even robotic vibrations scared crickets (Adamo et al., 2013), sounds emitted by bats can launch avoidance of moths (Cinel and Taylor, 2019), and real but harmless spiders with manipulated mouthparts can scare grasshoppers (Schmitz et al., 1997). Yet the effect of these stimuli can differ in the ecological context and among taxa (Cinel et al., 2020). Specifically, it remains unclear which signals the insect needs to receive to develop the stress response. Although birds represent one of the main predators of insects (Van Bael and Brawn, 2005; Bael et al., 2008), the knowledge about the stress response of insects to the presence of birds was not evaluated directly. We can only expect that insects sense the presence of birds (Fournier et al., 2013).

To fill this knowledge gap, we evaluated the hormonal and behavioural responses in desert locusts (*Schistocerca gregaria)* (Forskål, 1775), which were exposed to (1) live birds which represented a real, life-threatening risk, (2) warning calls of the birds which represented only a potentially threatening signal, and to (3) control conditions without the sign of the presence of predator. We conducted the experiments in indoor experimental test cages as well as in outdoor aviaries. We hypothesised that the acoustic signal, such as bird call, represents a clue of the predator presence which will lead to a detectable stress reaction. The presence of birds was then expected to posea real, life-threatening risk, after which we anticipated to find a strong stressful reaction, potentially stronger than after exposure of locusts to bird call only (Fournier et al., 2013).

## Methods

### Experimental animals

Desert locust, *Schistocerca gregaria* (Forsskål, 1775), occurs in Africa, the Middle East, and Asia (Topaz et al., 2012). The desert locust is polyphagous and feeds on leaves, shoots, flowers, fruit, seeds, stems, and soft bark, and is well known to periodically form enormous destructive swarms. At least 30 bird species were observed feeding on locusts, including 10 European wintering species (Sánchez-Zapata et al., 2007). The desert locusts perceive sound by tympanal organs in a frequency 0-20 kHz (Haskell, 1957), however its ability to perceive sound has been questioned (Gordon et al., 2014). The color vision system consists of green, blue and UV sensitive visual fibers (Schmeling et al., 2014) but it is not known to what distance they are able to see.

Vibrations are detected by mechanoreceptors in the legs, the scolopidial organs (Lakes-Harlan and Strauß, 2014).

For our experiments, the subadults (4^th^ instar, 3-4 cm long) of gregarious form desert locusts were purchased from a commercial supplier (Acheta.cz, Czech Republic). After purchase they were placed in a calm room with natural light, where they were either kept individually in small plastic containers for 48 hours (before experiment I) or together in a large cage, where food and water were provided *ad libitum* (before experiments II and III). None of the individuals had previous experience with any predators.

Great tit (*Parus major* L., 1758) was selected as a representative of insectivorous birds, which inhabits Europe, Northern Africa, and large areas in Asia. Although desert locusts are not typical prey of great tits in Czech Republic, they have been reported as their prey in migratory grounds where the desert locusts occur (Mullié, 2009). Further, interspecific eavesdropping on alarm calls, which are typically 3-5 kHz (Fallow et al., 2013), has been confirmed for many species, thus any bird alarm call can be expected to start a stress reaction in arthropods.

The great tits were mist-netted in the proximity of the Faculty of Science, University of South Bohemia a day or two prior to planned experiments with locusts and kept in accredited breeding areas. They were housed individually in cages with dimensions 0.7 × 0.4 × 0.5 m with a plastic bottom and three perches. The birds were provided with food (sunflower seeds, mealworms, insect patee for passerines) and water *ad libitum* daily. They were released immediately after the experiment, so they were in the breeding areas for up to 3 days. New birds were captured for the next trial.

All experiments were conducted in the laboratories of the Institute of Entomology, Biology Centre CAS, and in the aviaries of Faculty of Science, University of South Bohemia which have joined campus in Ceske Budejovice, Czech Republic from May to October 2018.

### Laboratory experiment I – behavioural test

The laboratory experiments were conducted in a quiet experimental room with natural light conditions and in modified bird cages with perforated plastic bottom part to enable air circulation. The cages had dimensions 0.7 × 0.4 × 0.5 m and their two parts were separated by 0.2 mm wide non-glossy glass. The bottom was aimed for locusts and the upper part of the cage for birds. Two independent experimental cages were located 2 m apart in the experimental room. The position of the cages in the room was noted (either left or right side of the room) to document its potential effect on the results.

Prior to the behavioural experiment, the locusts were starved for 3 days to keep them motivated for food searching. The shelter for locusts (a 15×10 cm part of the egg carton) was placed to the bottom of each cage. Food (two halves of a grape, a piece of carrot, and lettuce) on a Petri dish was on the opposite side of the cage.

The space between the shelter and the food, i.e., area of 0.3 × 0.3 m, was considered as an open field. Immediately prior to the experiment, five locusts were placed under the shelter in the bottom part of each cage.

Locusts were exposed to three different treatments: (1) “bird treatment” – one live great tit present in the cage plus a playback of a mixture of songs and warning calls of great tits (see below), (2) “call treatment” - the playback of the bird calls only, and (3) “control” – no bird and silence in the experimental room. The playback of the bird calls was a 60 min long mixed sequence of alarm calls from the Xeno-canto collection (www.xeno-canto.org). The playback was played using a JB.lab R1 Full-range speaker. To limit the effect of vibrations on the locust perception, the cages were set and tightened to the desk to avoid vibrations or movement of the cage itself. The experiments were run in the following order: control, call, bird treatment to avoid the potential effect of chemical cues released by locusts or birds. After the bird treatment the experimental room was aired for at least 2 hours. Then another set of sequences of the treatments was launched.

Their behaviour was recorded by two cameras (Sony SHD-CX240E) positioned on a tripod above the cage. The recording in the absence of an observer lasted for 60 minutes. The time spent in the shelter and time spent feeding was scored in seconds for individual locusts. We also noted whether the locusts entered the open field between the food and the shelter (sometimes, the locust just sat next to the shelter but did not move away from it towards the food). If the locusts have not left the shelter or have not reached the food, the time was scored as 3600 sec. The experimental locusts were discarded after each trial.

### Laboratory experiment II – physiological test

In the second stress induction experiment, we evaluated Schgr-AKH-II levels in the CNS and haemolymph of locusts under laboratory conditions. Schgr-AKH-II (pGlu-Leu-Asn-Phe-Ser-Thr-Gly-Trp-NH2) (Gäde et al., 1986) as one of two AKHs in *S. gregaria* was used as a marker. In contrast to the laboratory experiment I, the locusts were fed before the experiment *ad libitum* and 10 locusts were moved to the plastic bottom of each cage where neither food nor shelter was provided. The locusts were moved to the experimental cages just prior to the beginning of the experiment as quickly as possible and then exposed for 30 min (Orchard and Lange, 1983) to the three treatments.

### Outdoor experiment III – physiological test

To simulate more natural (i.e., fresh circulating air, realistic distances of prey from birds, higher number of communicating birds) experimental conditions, the outdoor aviary (10 × 15 m) with shrubs, small trees and high grass was established in the campus of Biology Centre CAS (48°58’37.1”N 14°26’49.2”E). A flock of 20 young great tits was released into the aviary one week prior to the experiment. The birds were receiving the food and water *ad libitum* at several places within the aviary. The bird and the control (birds absent) treatments were conducted, with 4 perforated bottom parts of the cage covered by the non-glossy glass with 10 locusts in each treatment. The glass prevents the direct attack of birds to locusts. The locusts in the bird treatment were placed on the ground of the aviary. The exposure of locusts to the bird treatment lasted for 3 hours (9-12 am), ambient temperature was 20-25 °C. The bottom parts of cages with locusts were naturally shaded by surrounding vegetation to avoid their overheating. The control treatment was deposited in the nearby shady area (48°58’36.7”N 14°26’45.5”E) with the absence of birds.

### Schgr-AKH-II extraction from CNS and haemolymph

Immediately after the finishing of the experiments II and III haemolymph was taken from the locust bodies (see below) and their heads were cut off under the Ringer saline. The brain with corpora cardiaca and corpora allata attached (= CNS) was dissected from the heads and AKH was extracted from the CNS using 80% methanol. The solution was evaporated in a vacuum centrifuge and the resulting pellet stored at -20°C until needed.

For determination of the endogenous Schgr-AKH-II titre in the haemolymph by ELISA, some pre-purification steps described in our previous paper (Goldsworthy et al., 2002) were essential. Briefly, haemolymph samples were collected from the locusts by a piercing of the soft cuticle between the hind leg and thorax.

After piercing, the leaking haemolymph was collected by pipette tips. In total, we obtained and pooled at least 300 μl of haemolymph from each group of 10 locusts per cage and treatment and collected it into 1.5ml Eppendorf tubes, both in laboratory and outdoor experiment. Then, the haemolymph was stored at -26°C. Later on, it was extracted in 80% methanol and after centrifugation the supernatants were evaporated to dryness. Then the dry pellets were dissolved in 0.11% trifluoroacetic acid, applied to a solid phase extraction cartridge Sep Pak C18 (Waters), and eluted by 60% acetonitrile. The eluent was analysed on a Waters HPLC system with a fluorescence detector Waters 2475 (wavelength λ_Ex_ – 280 nm; λ_Em_ – 348 nm) using a Chromolith Performance RP-18e column (Merck), solutions A and B (A – 0.11% trifluoroacetic acid in water; B – 0.1% trifluoroacetic acid in 60% acetonitrile) and a flow rate 2 ml/min. Fractions eluting between 13.2-14.4 min were subjected to competitive ELISA. Retention time of the *S. gregaria* AKH - Schgr-AKH-II was 13.8 min under the used conditions.

### ELISA determination of Schgr-AKH-II level

For determination of Schgr-AKH-II level in *S. gregaria* CNS and haemolymph a common direct ELISA was used. Thus, the ELISA comprised coating of the 96-well microtitre plate (high binding Costar, Corning Incorporated, Corning, NY, USA) by the extracts of one CNS or 40 μl haemolymph equivalent. The primary rabbit antibodies used in the procedure (dilution 1:1000) were raised commercially against the Schgr-AKH-II by Moravian-Biotechnology (Brno, Czech Republic); to exclude a possible cross-reactivity, the corresponding pre-immune serum was used as well. Swine anti rabbit IgG labelled with horseradish peroxidase (SwAR/HRP - LabNed) (dilution 1:2000) was used as a secondary antibody. And finally, the ELISA substrate 3,3’,5,5’- tetramethylbenzidine (Sigma Aldrich) was used to visualise the reaction. The absorbance values were determined in a microtiter plate reader at 450 nm.

### Statistical analysis

The behavioural response to the three treatments (Bird, Call, Control) was analysed as linear mixed effect models (lme) in the package nlme (Pinheiro et al., 2007) in R 4.0.2 (Team, 2020). The time spent in shelter (in sec) and time spent feeding (in sec) were explained by the treatment (*N* = 3) and position of the cage (i.e., cage identity; *N* = 2 in each of the 3 trials consisting of three treatments), while the identity of locust was used as a random effect. Tukey post-hoc test from package multcomp (Hothorn et al., 2015) was used to compare all possible pairs of mean values of the time spent in shelter and of the feeding time among the individual treatments.

Significance of differences among the treatments in the levels of the stress hormone in haemolymph and in CNS were tested by one-way ANOVA using GraphPad Prism 6.0 (GraphPad Software, San Diego, CA, USA). The effect of three (bird, call, control) and two treatments (bird, control) was tested separately for the laboratory and outside aviary experiments, respectively. The ANOVA statistics were followed by Tukey’s multiple comparison test (laboratory experiments) resp. by Student’s *t*-test (outdoor experiment) to compare effects of treatments.

### Ethical note

This research was conducted under the ethical approval of the University of South Bohemia. The wild *P. major* individuals were captured and held under the permit CZ5031 and permit of the city council of Ceske Budejovice, Czech Republic. The experiment on animals were conducted under 1415-20424/2019-65 permit issued by the Ministry of Environment, Czech Republic.

## Results

### Laboratory experiment I – behavioural test

Locusts spent a significantly longer time hiding when in the presence of a bird than in the call and control treatments (Fig. 1a; *F*_2,155_ = 35.994, *P* < 0.0001). In the presence of a bird, the locusts hid for approximately 2045.3 (± 115 SE) seconds. In contrast, in the call and control treatments, the locusts hid for 1102.5 (±102 SE) and 931.4 (± 111 SE) seconds, respectively. Cage position did not have an effect on the duration of hiding (*F*_2,155_ = 1.744, *P* = 0.178).

**Figure 1.**
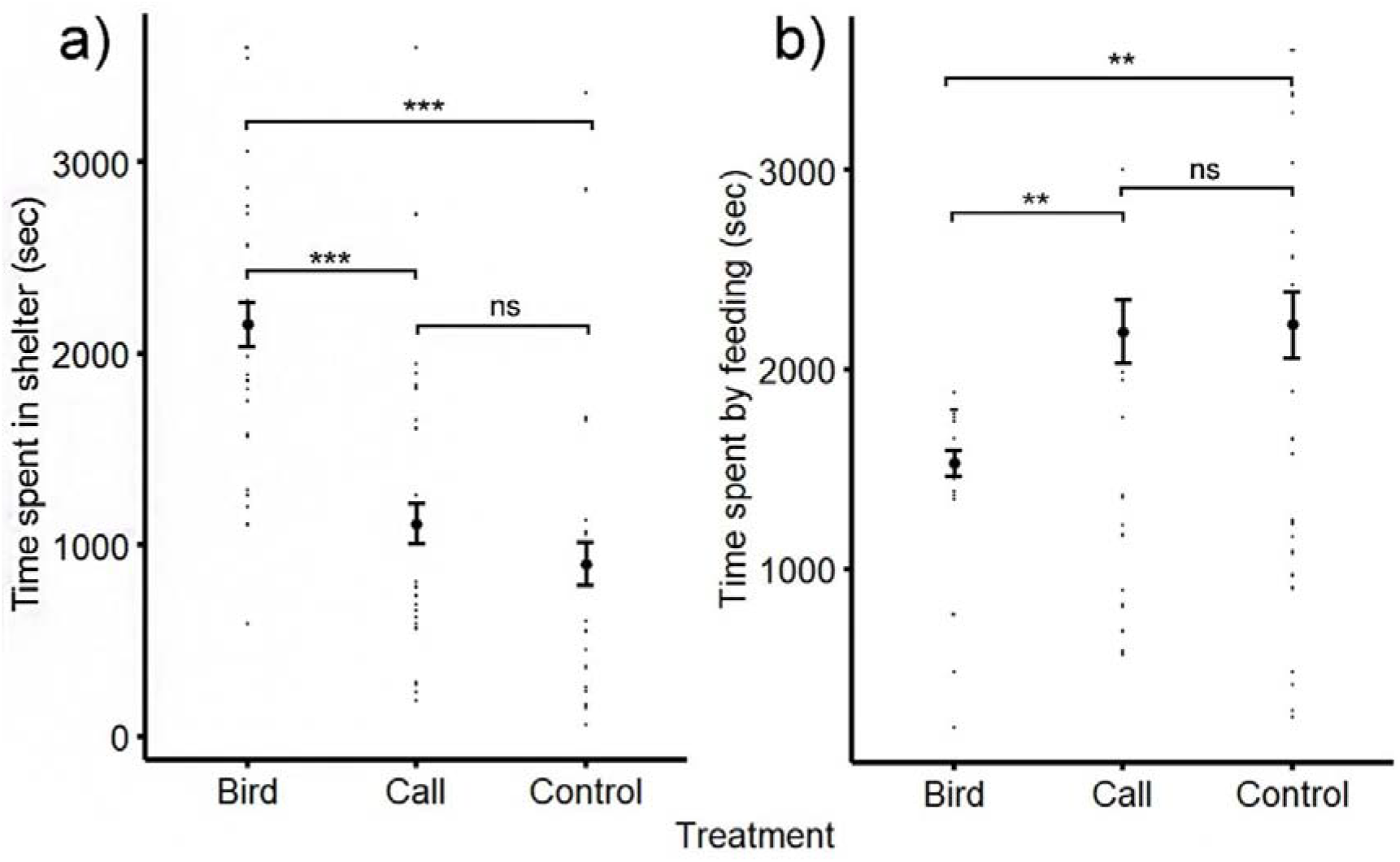
Effect of the bird (bird + bird call), the call (bird call only) and the control (neither bird nor call) treatments on the behaviour on adults of the *Schistocerca gregaria* locusts. Time spent in the shelter (a) and time spent feeding (b) in seconds is presented on axes-y. The significant results are marked by *** *P* < 0.001, ** *P* < 0.01, ns *P* > 0.05. Small dots represent raw values for the individual locusts and full points and whiskers show means ± SE (seconds) derived from the respective minimal models.

Congruently, time spent feeding was affected by the treatment (Fig. 1b; *F* _2,155_ = 7.406, *P* = 0.0008). The locusts were feeding in average 1530 (± 65 SE) seconds in the bird treatment, but only 2190 (± 162 SE) and 2223 (± 168 SE) seconds in the call and control treatment, respectively. Again, the cage position did not influence feeding time (*F* _2,155_ = 1.349, *P* = 0.263).

### Laboratory experiment II – physiological test

The presence of live birds and their calls in (i.e., bird treatment) induced a significant increase in the Schgr-AKH-II in both the CNS (1.4 times, one-way ANOVA: *F* = 5.725, *P* = 0.0056, and haemolymph (1.3 times, one-way-ANOVA: *F* = 32.12, *P* = 0.0001) of the locusts (Fig. 2). Interestingly, application of the bird call only (call treatment) had no significant effect on either the CNS (Tukey’s posttest: *P* = 0.9835) or haemolymph (Tukey’s posttest: *P* = 0.2375).

**Figure 2.**
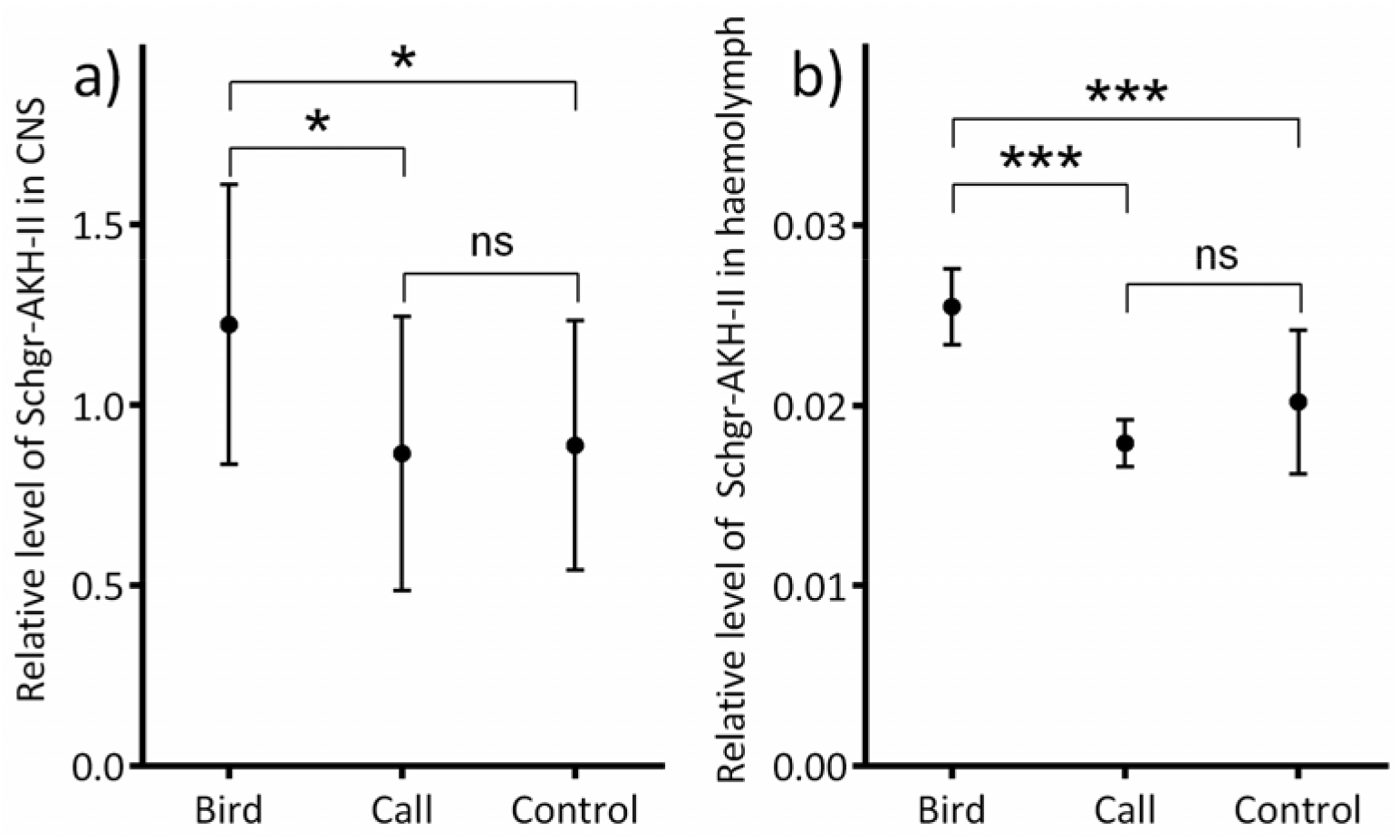
Effect of predator treatments on the Schgr-AKH-II level in CNS (a) and in haemolymph (b) of *S. gregaria* within the laboratory experiments. Statistically differences between the groups were evaluated using one-way ANOVA with Tukey’s posttest, and significant results are marked by *** *P*<0.001, * *P*<0.05, ns *P*>0.05. Treatments: Bird = real great tits were present in the experimental cages and warning calls were played, Call = only warning call of great tit was played, Control = only locusts were present in the cages.

### Outdoor experiment III – physiological tests

The presence of live birds and their calls (bird treatment) as a stress factor induced a significant increase in the Schgr-AKH-II level in both the CNS (1.4 times, one-way ANOVA: *F* = 5.725, *P* = 0.0056) and haemolymph (1.3 times, one-way-ANOVA: *F* = 32.12, *P* = 0.0001) of the locusts (Fig. 3). Interestingly, application of the bird call only (call treatment) had no significant effect on either the CNS (Tukey’s posttest: *P* = 0.9835) or haemolymph (Tukey’s posttest: *P* = 0.2375).

**Figure 3.**
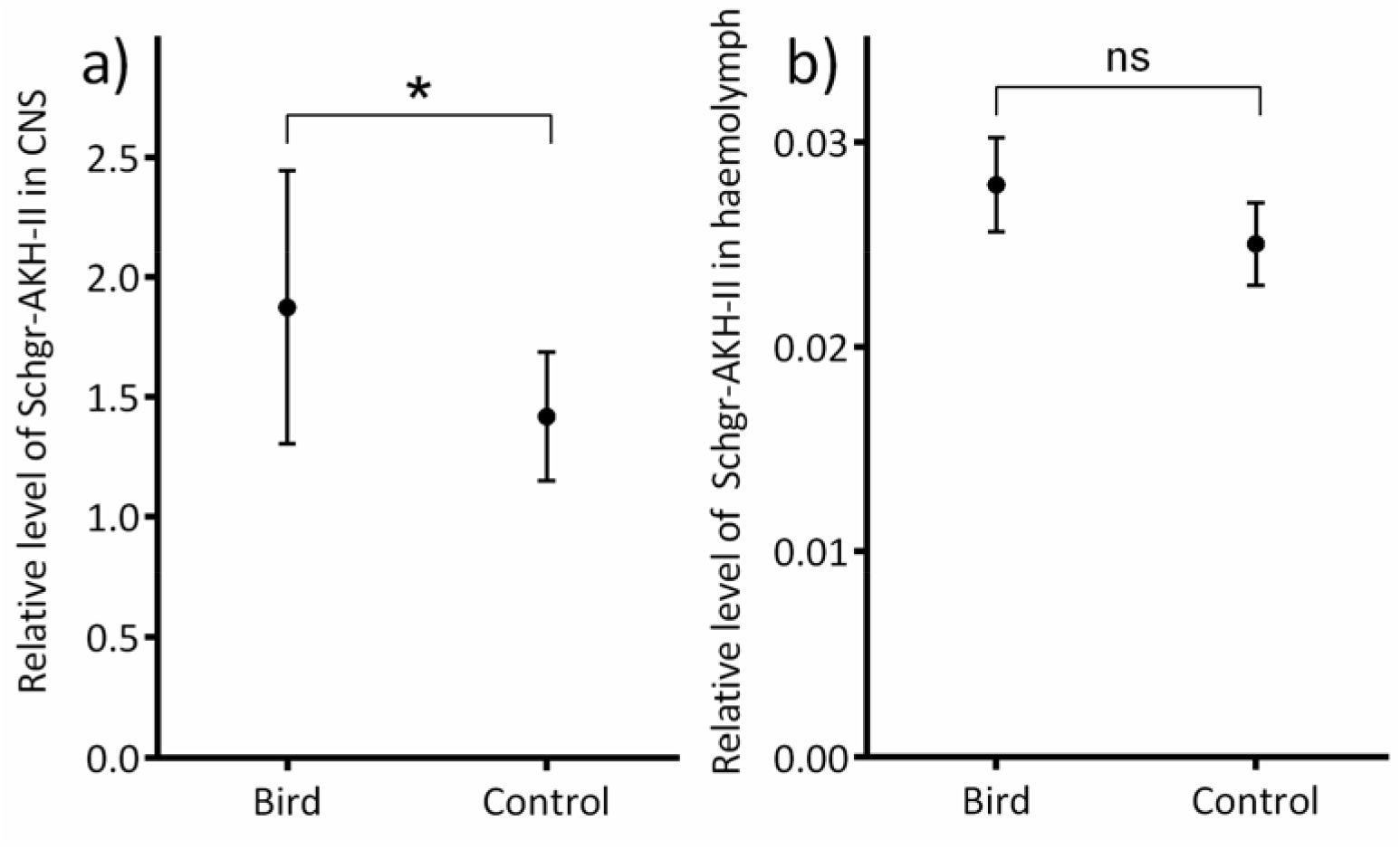
Effect of predator treatments on the Schgr-AKH-II level in CNS (a) and in haemolymph (b) of *S. gregaria* in the outdoor aviary experiments. Statistically significant differences between the groups were evaluated using the Student’s *t*-test, and significant results are marked by * *P* < 0.05, ns *P* > 0.05 (*n* = 4-16). Treatments: Bird = real great tits were present in the aviary and warning calls were played, Control = only locusts were present in the nearby shady area without birds.

## Discussion

We found that the presence of the great tits drives a hormonal as well as behavioural response in gregarious desert locusts. The desert locusts spent twice as much time under the shelter and limited the feeding time by 60% when a real bird and its alarm call was presented to them, compared to controls. Levels of stress hormones in the CNS and haemolymph were significantly higher in the presence of the real bird compared to the bird call and control treatments. This provides novel information which integrates the knowledge about interaction among the level of stress hormones and behaviour in insects. It is also in line with our expectations and would mean that fear of birds might result in lower feeding rate, slower growth and maybe lower reproduction thus shaping evolution of interactions between insects and plants (Fournier et al., 2013).

Against expectations (Minoli et al., 2012) the alarm call of great tits was not an appropriate signal to induce hormonal and behavioural change in gregarious desert locusts. The time devoted to hiding and feeding during the continuous alarm calling of great tits was not significantly different from the control treatments. Several alternative explanations are possible. The bird singing and calling are nearly ubiquitous in nature and the cost of hiding permanently would be high. It is known that stress hormones can reduce predation (Adamo et al., 2013), but a permanently occurring level of stress hormones could be debilitating to the animal (Clinchy et al., 2013) It might be also speculated that a warning call is usually used by birds in response to bird’s predators, which might mean that the bird is less likely to continue the search for prey and can be evaluated as less dangerous by the insect. Unlike bat vocalisation used to locate prey, the bird calls may not represent an immediate risk of attack (Cinel and Taylor, 2019). Further, it might be because desert locusts did not perceive the great tit as a potentially risky predator, but they were afraid of silhouette or movement as birds are generally predators of locusts. This would however suggest that locusts recognize the specific call of great tit but generalise the silhouette.

Final explanation might lay in the locust’s ability to hear. Locusts have a well-developed auditory system that is able to determine a wide range of pitch. Desert locusts show extreme phenotypic plasticity forming, solitary and gregarious phases, which differ extensively in behaviour, physiology including their sensory abilities, and morphology (Simpson et al., 1999; Gordon et al., 2014). In previous experiments, gregarious forms of desert locusts responded in 6 out of 12 cases to swarm sounds by an escape (Haskell, 1957) and took evasive action after hearing ultrasound calls of hunting bats (Weber et al., 1981). Yet, Gordon et al (2014) found differences in audition of both forms of locusts. Solitarious locusts fly at night (Ould Ely et al., 2011), and hence are potentially at much greater risk from predation by bats b(Haskell, 1957; Robert, 1989) which seems to be in line with their sensitivity to the higher frequencies used by bats in their echolocation calls (Gordon et al., 2014). On the other hand, gregarious locusts are more active during the day and so they should need a stronger detection of birds. However, our current observation that the desert locusts did not react to bird alarm calls cannot support this hypothesis. Future work could test whether the locust responds to other, more appropriate, calls of predators.

Existing studies have revealed that various arthropods have variable responses to different types of predator-induced stimuli. Lohrey et al. (2009) found that spiders were freezing when a seismic or acoustic stimulus was presented, and they increased movement when they were exposed to a visual stimulus. Desert grass spider *Agelenopsis aperta* exhibited antipredator behaviour to puffs of air simulating bird wing beats (Riechert and Hedrick, 1990). Some crickets were shown to respond to species-specific vibrations that lizards made when walking (Adamo et al., 2013). Even placement of a dummy predator (robotic hamster) to terrarium resulted in changes in behaviour of the crickets (Adamo et al., 2013). In another experiment, the mere presence of spiders with their mouths glued shut changed the behaviour of the grasshoppers (similarly to predation treatment, where the spiders were allowed to eat them), which resulted in the grasshoppers acquiring less food, which in turn decreased grasshopper populations (Schmitz et al., 1997). The indirect effects of insectivorous arthropods on arthropods have been further proved through experiments in a number of systems: mantids and their predation on herbivorous insects (Moran and Hurd, 1997), effect of large predatory mosquitoes on smaller mosquitoes (Chandrasegaran and Juliano, 2019), effect of Anolis lizards on Homoptera and Araneae (Spiller and Schoener, 1990a; b), effect of beetle larvae on ants (Letourneau and Dyer, 1998), effect of sound of wasps on Lepidoptera larvae (Lee et al., 2021). Orthoptera specifically were shown to be able to detect risk by the air particle movement generated by the predator wing beat due to the filiform hair sensilla (Gnatzy and Kämper, 1990).

However, birds have been rarely used as the source of fear in terrestrial experiments with insects, although they are one of their main predators. Signals of bird presence are visual (whole animal, part of an animal or just its shadow), localized disturbance (e. g. vibration, movement of leaves, air movement) or different types of vocalizations. The perceived signals might be dependent on the distance and duration in which the arthropod and predatory bird interact. The dimensions of our birdcage in the indoor conditions were only 0.7 × 0.4 × 0.5 m, so the laboratory experiment was intense, as the locusts were in close contact with the bird, and they could see its movement. In contrast, the locusts in outdoor aviaries were not in close contact with the birds, as the birds typically moved 1 - 2 m far from the locusts. Pitt (1999) used 5 × 5 m large aviaries protecting grasshoppers from birds and observed that grasshoppers were significantly more often lower in vegetation but did not change their mobility and feeding behaviour. In other avian-free treatments, free living birds were not able to access the locusts closer than 2 metres, while they had free access to the locusts in the control treatment. The activity of locusts tended to be lower in the control plots; however, the pattern varied over the summer season. Similarly, to earlier study, the locusts tended to forage deeper in grass and did not climb so high, when birds were present (Belovsky et al., 2011).

Another crucial feature in the experiment could be the duration of the exposure of the prey to predators. Our 30-minute laboratory experiment (with a predator being close) and 3 hours long outdoor experiment (with a predator) resulted in similar levels of stress hormones. In general, a 30 min long exposure to predators was sufficient to detect the hormonal stress response. This would be in line with an earlier study, which found that the concentration of AKH in the haemolymph of *S. gregaria* changes and increases with the time of exposure to the stress factor but appears as quickly as in 2 minutes and peaks after 60 minutes (Candy, 2002).

The link between various components of the stress response system and behaviour controlled by the nervous and endocrine systems is far from a proper understanding (Storey, 2004; Johnstone et al., 2012). Nevertheless, in arthropods, biogenic amines and certain neurohormones seem to play a crucial role in control of stress response (Kodrik, 2008; Nelson et al., 2021). It is obvious that these amines are involved directly into the defence reactions, e.g. octopamine modulates anti-predator behaviour in beetles (*Tribolium castaneum*) (Nishi et al., 2010) and in an orb-weaving spider (Jones et al., 2011). On the other hand, biogenic amines are thought to be involved in regulation of AKH production (Van der Horst et al., 2001) or the AKH release from the corpora cardiaca into haemolymph (Orchard et al., 1993). There are many examples in the literature describing fluctuations in AKH levels in the insect CNS and haemolymph after a stressor application. For example, 20 minutes of forced running (in a horizontal laboratory shaker) resulted in a slight increase of AKH titre in the CNS of the firebug, *Pyrrhocoris apterus*, but in a strong increase of the titre in the haemolymph (Kodrík and Socha, 2005). A similar picture can be seen when applying various pathogens (Ibrahim et al., 2017; Ibrahim et al., 2018; Gautam et al., 2020a; Gautam et al., 2020b) or toxins including insecticides (Candy, 2002; Kodrík et al., 2015). Changes in the AKH levels in the haemolymph are often faster, while in the CNS are slower but more profound. Nevertheless, in this study, all changes in Schgr-AKH-II titre after the bird treatment were similar except for the no change of Schgr-AKH-II in the haemolymph within the external experiment. To explain this, we can only speculate that due to the large distance of the locusts from the birds, the stress was not strong enough, as mentioned above.

In conclusion, we report evidence that insects in the presence of birds decreased their foraging activity, reducing their food income roughly by 60%, and we relate the results of behavioural tests to a physiological stress response. Our experimental results are in line with the risk allocation hypothesis (Lima and Dill, 1990; Lima and Bednekoff, 1999) predicting that animals should increase their foraging effort in the low-risk environment and decrease foraging in the high-risk environments. We argue that the methodological approach combining behavioural experimentation with physiological assessment of animal responses should be crucial for insight into fear perception in animals and for understanding their fitness optimization strategies. Last, future tests should focus on the effect of bird calls on behaviour of other insect species to extend the information gained by our study.

## Authors contributions

Jan Kollross – conceptualization, experimental investigation, writing of the first draft, visualization; Jitka Jancuchova-Laskova-experimental investigation, writing of the first draft together with JK; Irena Kleckova – methodology, experimental investigation; Inga Freiberga - experimental investigation, ; Dalibor Kodrik – conceptualization, laboratory analyses of the physiological responses; Katerina Sam - supervision, conceptualization, methodology, formal analysis, resources & funding acquisition. All authors contributed equally to editing of the manuscript.

## Acknowledgements

We acknowledge Matthias Weiss and Jakub Rehula for help with the experiments. The project was financially supported by ERC Starting Grant BABE 805189. Jakub Rehula was supported by project Open Science for grammar school students.

